# SGLT2 inhibition by intraperitoneal dapagliflozin mitigates peritoneal fibrosis and ultrafiltration failure in a mouse model of chronic peritoneal exposure to high-glucose dialysate

**DOI:** 10.1101/2020.11.04.366724

**Authors:** Michael S. Balzer, Song Rong, Johannes Nordlohne, Jan D. Zemtsovski, Sonja Schmidt, Britta Stapel, Maria Bartosova, Sibylle von Vietinghoff, Hermann Haller, Claus P. Schmitt, Nelli Shushakova

## Abstract

Peritoneal dialysis (PD) is limited by glucose-mediated peritoneal membrane (PM) fibrosis, angiogenesis and ultrafiltration failure. Influencing PM integrity by pharmacologically targeting sodium-dependent glucose transporter (SGLT)-mediated glucose uptake has not been studied. In this study wildtype C57Bl/6N mice were treated with high-glucose dialysate via an intraperitoneal catheter, with or without addition of selective SGLT2 inhibitor dapagliflozin. PM structural changes, ultrafiltration capacity and PET status for glucose, urea and creatinine were analyzed. Expression of SGLT and GLUT was analyzed by real-time PCR, immunofluorescence and immunohistochemistry. Peritoneal effluents were analyzed for cellular and cytokine composition. We found that peritoneal SGLT2 was expressed in mesothelial cells and in skeletal muscle. Dapagliflozin significantly reduced effluent TGF-β concentrations, peritoneal thickening and fibrosis as well as microvessel density, resulting in improved ultrafiltration, despite the fact that it did not affect development of high glucose transporter status. *In vitro*, dapagliflozin reduced monocyte chemoattractant protein-1 release under high glucose conditions in human and murine peritoneal mesothelial cells. Pro-inflammatory cytokine release in macrophages was reduced only when cultured in high glucose conditions with an additional inflammatory stimulus. In summary, dapagliflozin improved structural and functional peritoneal health in the context of high glucose PD.

## INTRODUCTION

Peritoneal dialysis (PD) as a renal replacement therapy for individuals with end stage renal disease relies on the peritoneum and its properties as a dialyzer membrane. Glucose-based PD fluid (PDF) generates an osmotic gradient that promotes water and solute clearance across the peritoneal membrane. However, glucose-containing PDF is non-physiological and as a result, in most PD patients structural and functional changes occur over time, resulting in decreased dialysis efficiency and ultimately technique failure. [1] While our understanding of the molecular mechanisms of such PD-related structural and functional aberrations of the peritoneum has grown considerably over the last decades, successful translation of pathophysiological insights into therapeutic options for peritoneal fibrosis are scarce.[2]

High glucose concentrations applied in PD create a diabetic state of the peritoneal cavity.[3] Mesothelial cells (MC) are the first cells of the peritoneal membrane that get in contact with glucose-containing PDF. The glucotoxic milieu itself can trigger detrimental changes in mesothelial cells such as epithelial-to-mesenchymal transition (EMT) and increased production of pro-inflammatory, pro-fibrotic and pro-angiogenic mediators promoting leukocyte infiltration, fibrosis and angiogenesis.[4] Although the detrimental effects of glucose uptake from the peritoneal cavity have received considerable attention in PD research,[5] studies on glucose transporters at the mesothelial cell level and their morphological and functional impact in the setting of PD are scarce. Several decades ago, studies demonstrated expression of sodium-dependent glucose transporter (SGLT)1 at the apical plasma membrane of human peritoneal mesothelial cells (HPMC).[6] Only recently, the existence of both SGLT1 and SGLT2 in the peritoneum has been demonstrated in rats. [7] Given the wealth of recent studies that implicate SGLT2 inhibition with antifibrotic properties not only in the kidney[8] but also in other organs such as liver[9] and heart,[10] we asked whether or not SGLT would be a feasible pharmacological target in PD patients in order to ameliorate structural and functional changes in the peritoneum.

To this end, we first confirmed the peritoneal expression of SGLT in mice and in human peritoneal biopsies. We then intraperitoneally applied the SGLT2 inhibitor dapagliflozin via a PD catheter-based chronic PDF exposure model to mice and evaluated its effects on peritoneal structure and function. We show that treatment with dapagliflozin ameliorated fibrotic and angiogenetic changes as well as ultrafiltration failure.

## MATERIALS AND METHODS

### Human peritoneal samples

Human peritoneal biopsies were biopsies taken from PD patients and non-uremic control patients undergoing surgery because of non-renal causes (excluding trauma, intra-abdominal neoplasia or inflammation) after informed consent according to the declaration of Helsinki and local ethics board approval at the Hannover (MHH #17/6715) and Heidelberg (S-493/2018) study sites. Peritoneal biopsies were processed and analyzed as described previously.[11,12] The non-CKD patient was 3 years old and underwent surgery because of reflux, had normal biochemical findings and no signs of inflammation. The PD sample was obtained from a 14 years old child with nephronophthisis who had been treated with Balance^®^ (Baxter) for 12 months.

### Peritoneal dialysis fluid exposure model in mice

All animal experiments were approved by the animal protection committee of the local authorities (Lower Saxony state department for food safety and animal welfare, LAVES, approval: 33.19-42502-04-16/2266). 12 weeks old female C57Bl/6N mice (Charles River) were subjected to chronic peritoneal dialysis fluid exposure as described previously.[4] In short, 2.0 mL of standard PDF composed of 4.25% glucose and buffered with lactate (CAPD/DPCA3, Stay Safe; Fresenius) or 0.9% saline solution for controls were instilled daily via a peritoneal catheter connected to an implanted subcutaneous mini access port (Access Technologies) for 5 weeks (n=5 saline, n=12 PDF). Dapagliflozin at a concentration yielding a dose of 1 mg/kg body weight was added to saline (n=6) and PDF (n=12), respectively. Because dapagliflozin is easily soluble in aqueous solution, no vehicle was necessary. On the last day of experiments functional analysis of the PM was performed by ultrafiltration and equilibration test and peritoneal effluents were sampled as previously described.[4,13] Thereafter, tissue samples were collected from the anterior abdominal wall for histological and immunofluorescence analysis.

### Chemical analyses of blood and urine, peritoneal ultrafiltration and transport studies

24h urine collections, dialysate effluents and plasma were analyzed for glucose, creatinine, and urea using an Olympus AU480 chemistry analyzer. 2.5 mL of PDF was instilled into the peritoneal cavity and the mice were sacrificed after 120 minutes. The total intraabdominal peritoneal fluid was collected, and the drained volume was measured. Peritoneal ultrafiltration capacity was determined by the amount of peritoneal fluid recovered after 120 minutes. Recovered effluent was either analyzed immediately with flow cytometry or stored at −80 °C for further ELISA or biochemical analysis.

As surrogates for peritoneal solute transport at time point 120 minutes, we calculated dialysate-to-dialysate_0_ (D/D_0_) for glucose as well as dialysate-to-plasma (D/P) ratios for creatinine and urea. The transport of small solute was also evaluated by the mass transfer area coefficient (MTAC), using the Garred two-sample model:[14]

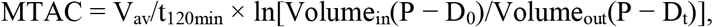

in which V_av_ is the average of the initial and final volumes; P, the plasma concentration of urea; D_t_, the dialysate concentration of urea or creatinine at the end of the dwell; D_0_, the initial concentration of urea or creatinine in dialysate, which is set at 0.

### Flow cytometry and ELISA measurements in peritoneal effluents

The inflammatory cell populations in the effluents were analyzed by flow cytometry using a FACS Canto II cytometer (BD Biosciences). The following monoclonal antibodies (BioLegend) were used: anti-CD11b (clone M1/70), anti-F4/80 (clone BM8), anti-CD19 (clone 6D5); anti-Gr1(clone RB6-8C5), and anti-TCRb (clone H57-597). Data were analyzed using FlowJo software (Tree Star). Inflammatory cytokines IL-6, IL-10, IFNγ, TNF-α and MCP-1 were analyzed by bead-based flow cytometry assay (CBA kit, BD Biosciences), TGF-β and VEGF-A with specific ELISA (R&D Systems) according to the manufacturer’s instructions.

### Morphological, immunofluorescence and immunohistochemical analysis of peritoneum

Submesothelial thickness of the peritoneum was determined on 2.5 μm paraffin-embedded tissue sections stained with Masson’s trichrome (Sigma-Aldrich) by blinded microscopy analysis (DM-IL microscope, DC300F camera, IM500 software, all Leica Microsystems). To allow for an unbiased analysis, thickness values were expressed as the mean of 40 independent measurements per animal at standardized interspaced locations of the peritoneum. Collagen I and III positivity was analyzed on sections stained with picrosirius red (Sigma-Aldrich) using an integrated intensity thresholding method detailed in Supplementary Methods (ImageJ software); results are given as percentage of total tissue area. Tissue sections were stained with primary antibodies against SGLT1 (Millipore 07-1417), SGLT2 (Abcam ab85626) and CD31 (Dianova DIA310) respectively. Background control staining was performed by incubating with secondary antibodies alone, omitting the first antibodies, and proved to be negative. Cell nuclei were stained with DAPI or hematoxylin Harris. For automated microvessel imaging NanoZoomer 2.0-HT Scan System (Hamamatsu Photonics) was used at 20x magnification (resolution: 0.46 μm/pixel). The slide scanner automatically detects the region of interest (ROI) containing the tissue and automatically determines a valid focal plane for scanning. As PDF penetration level reaches 400 μm, an area reaching 400 μm below mesothelial cell layer was annotated as ROI and microvessel density was quantified by microvessel algorithm v1 (Aperio Image Scope, Leica).

### RNA extraction and real-time quantitative PCR

Total RNA was extracted from harvested anterior peritoneal walls not affected by the peritoneal catheter using RNeasy mini kit (Qiagen) and reverse-transcribed using Promega kits. Real-time quantitative PCR analysis was performed on a LightCycler480 (Roche) real-time PCR system using SybrGreen as well as TaqMan technologies; β-tubulin and Rn18S mRNA were used as reference genes. Quantification was conducted using the delta-delta Ct method.

### HPMC, immortalized MPMC and RAW264.7 macrophage cell culture and treatment

For *in vitro* experiments, 3 different cell types were analyzed: primary human peritoneal mesothelial cells (HPMC), immortalized mouse peritoneal cells (MPMC) and murine peritoneal macrophage cell line RAW264.7.

HPMC were derived from omentum samples of 3 human controls as described previously[4] and grown to 80% confluence. In short, HPMC were isolated with trypsin/EDTA digestion method from omentum tissue obtained from patients with normal kidney function undergoing elective abdominal surgery. Informed consent was obtained for the use of omentum tissue and the study was approved by the institutional ethics committee (Hannover Medical School #17/6715). The cells were grown in RPMI1640 medium supplemented with 10% fetal bovine serum (FBS), 100 U/ml penicillin and 100 mg/ml streptomycin

Immortalized MPMC were generated in our lab and cultivated as described previously[4]. Briefly, the cells were grown to 80% confluence in RPMI1640 medium containing 1% penicillin–streptomycin, 10% fetal calf serum, 1% insulin/transferrin/selenium A (all from Life Technologies, Carlsbad, CA), 0.4 mg/ml hydrocortisone (Sigma-Aldrich), and 10 U/ml recombinant mouse interferon-□ (Cell Sciences, Canton, MA) at 33 °C (permissive conditions) to 80% confluence. For experiments the cells were differentiated for 3 days in the same medium without interferon-□ at 37 °C (non-permissive conditions).

Murine macrophage RAW264.7 cells were grown to 80% confluence in RPMI1640 medium supplemented with 10% fetal bovine serum (FBS), 100 U/ml penicillin and 100 mg/ml streptomycin

All cells were starved overnight in 1% FCS–containing RPMI 1640 medium and then cultured in the medium with normal (10 mM, NG, control) or high glucose (120 mM, HG) concentration for 24 or 48 h. For inhibition of SGLT2, different concentrations of dapagliflozin ranging from 3 to 300 μM were added to culture medium. Thereafter, SGLT1 and SGLT2 expression was analyzed in MPMC by RT-PCR as described for mouse peritoneum and intracellular glucose concentration was measured in MPMC and RAW264.7 cell lysates using Olympus AU480 multianalyzer.

In some experiments MPMC and RAW264.7 macrophages were cultured under either NG or HG conditions with or without addition of dapagliflozin for 48 h, followed by additional stimulation with LPS (10 ng/mL) for 8h. Conditioned cell culture medium was then analyzed for MCP-1, TNF-α and IL-6 by bead-based flow cytometry assay (CBA kit, BD Biosciences).

### Statistical analysis

Data are presented as means ± SEM, if not stated otherwise. D’Agostino & Pearson omnibus normality test was used to test for normality. Multiple comparisons were analyzed by one-way ANOVA with Sidak’s *post hoc* correction or Kruskal-Wallis nonparametric test with Dunn’s *post hoc* correction. All tests were two-tailed. P<0.05 was considered to indicate statistically significant differences. GraphPad Prism 7 was used for data analysis.

## KEY RESOURCES TABLE

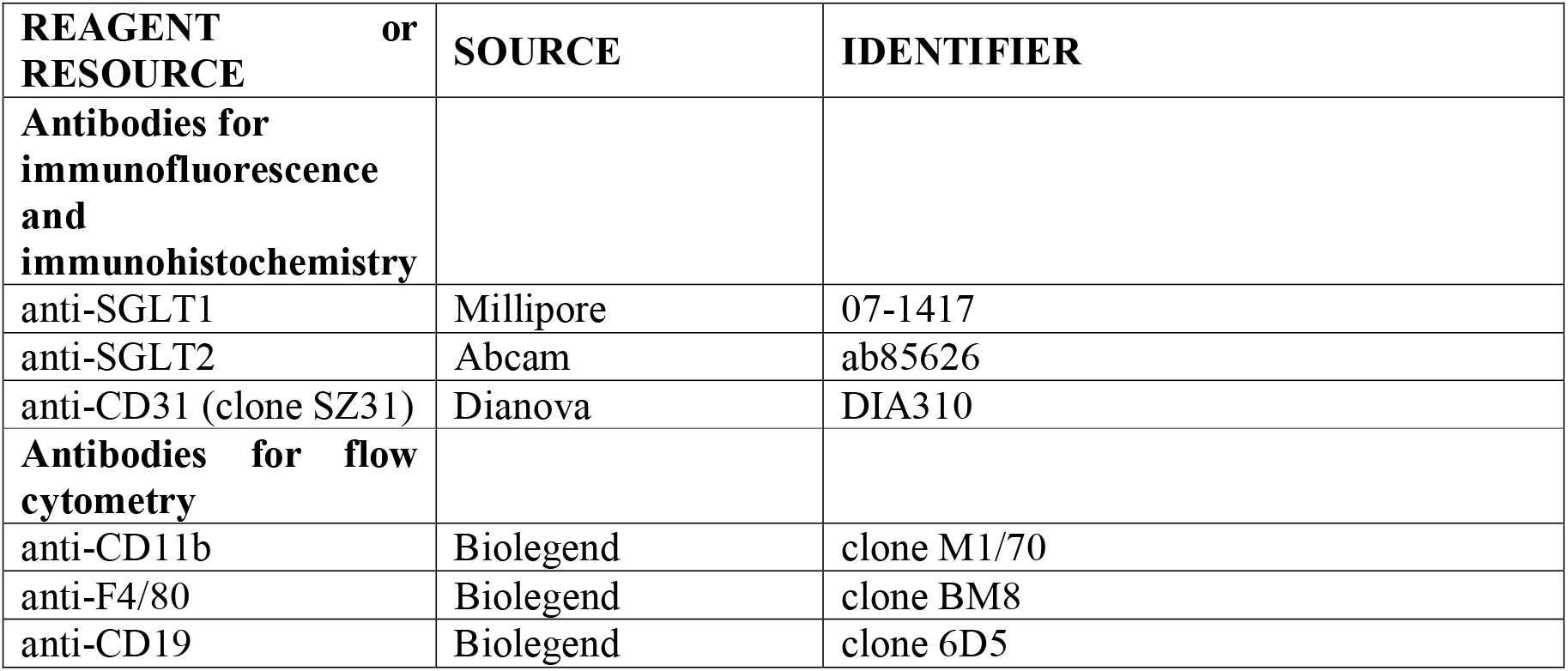
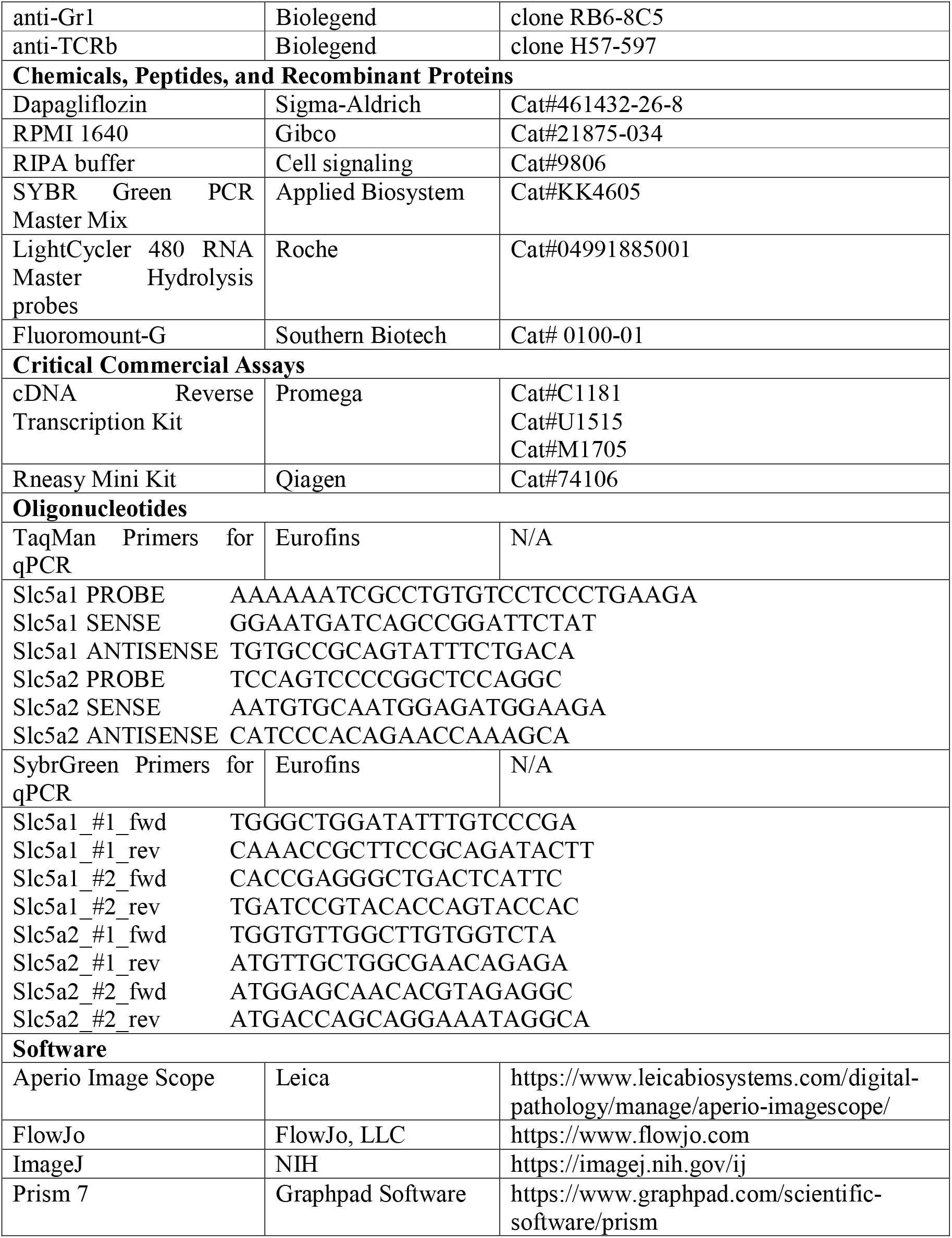

## RESULTS

### Sodium-dependent glucose transporters are expressed in the murine and human peritoneal membrane

First, we studied the presence of sodium-dependent glucose transporters in the peritoneal membrane. Using immunofluorescence, we demonstrate in 16 weeks old female C57BL6 mice that both SGLT1 and SGLT2 protein are expressed in the peritoneum, most prominently in the single mesothelial cell layer, but also in submesothelial skeletal muscle (**Figure 1a, upper row**). Antibody specificity against SGLT1 and SGLT2, respectively, was confirmed in kidney tissue from the same animals (**Figure 1a, lower row**). Moreover, using immunohistochemistry and immunofluorescence, we demonstrate presence of SGLT1 and SGLT2 in the human peritoneum in biopsies from healthy non-CKD control and PD patients, respectively (**Figure 1b**). In addition to the mesothelial cell layer, SGLT1 protein was visualized around capillaries in the submesothelial zone.

**Figure 1:**
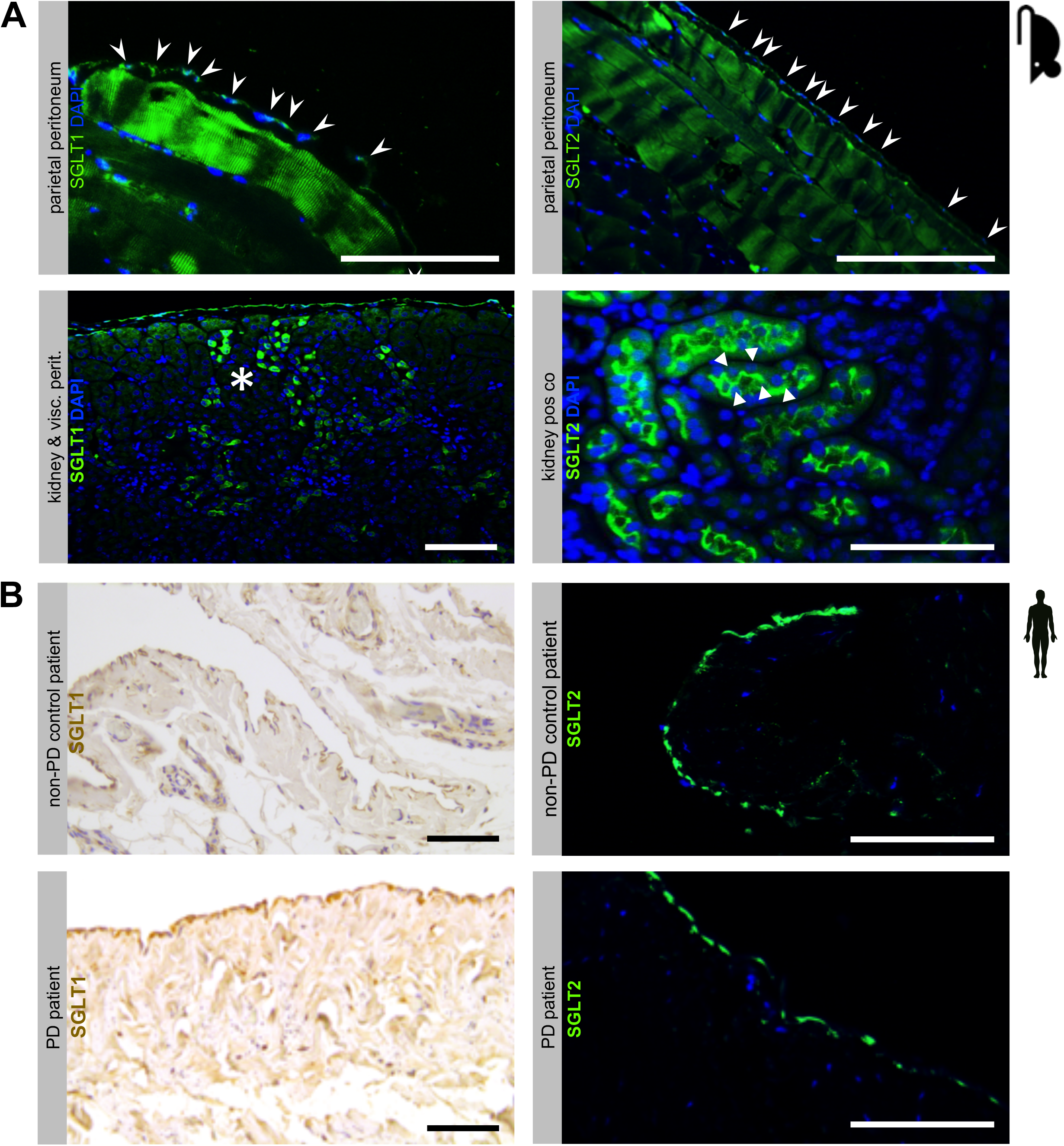
Expression of sodium-dependent glucose transporters (SGLT) at the murine and human peritoneal membrane. (A) Immunofluorescence staining of SGLT1 and SGLT2 in mouse peritoneal membranes. Antibody specificity is demonstrated in kidney positive controls, which show specific and distinct staining patterns of the proximal tubule brush-border membrane for SGLT1 (asterisk) and SGLT2 (arrowheads), respectively. Staining of the mesothelial cell layer for SGLT1 and SGLT2, respectively, is denoted by arrows. Blue staining denotes DAPI, scale bar=100 μm. (B) Immunohistochemistry and immunofluorescence staining for SGLT1 (left) and SGLT2 (right), respectively, in human peritoneal biopsies from non-PD control and PD patients, respectively. Note staining of the mesothelial cell layer and in the pericapillary region. Blue staining denotes DAPI, scale bar=100 μm.

### Chronic PDF-induced SGLT2 upregulation is abrogated by intraperitoneal dapagliflozin treatment

Next, we evaluated the influence of chronic glucose exposure on the peritoneal expression of sodium-dependent and sodium-independent glucose transporters and to analyze potential effects of pharmacological inhibition of SGLT2. To this end, we used the well-established mouse model of catheter-delivered chronic PDF exposure.[4] Mice were treated for 5 weeks with either saline or PDF with or without addition of dapagliflozin (1 mg/kg) via a peritoneal catheter (**Figure 2a**). Systemic action of dapagliflozin was observed, as reflected by presence of glucosuria in 24h urine collections of mice treated with the SGLT2 inhibitor (**Supplementary Figure 1**).

**Figure 2:**
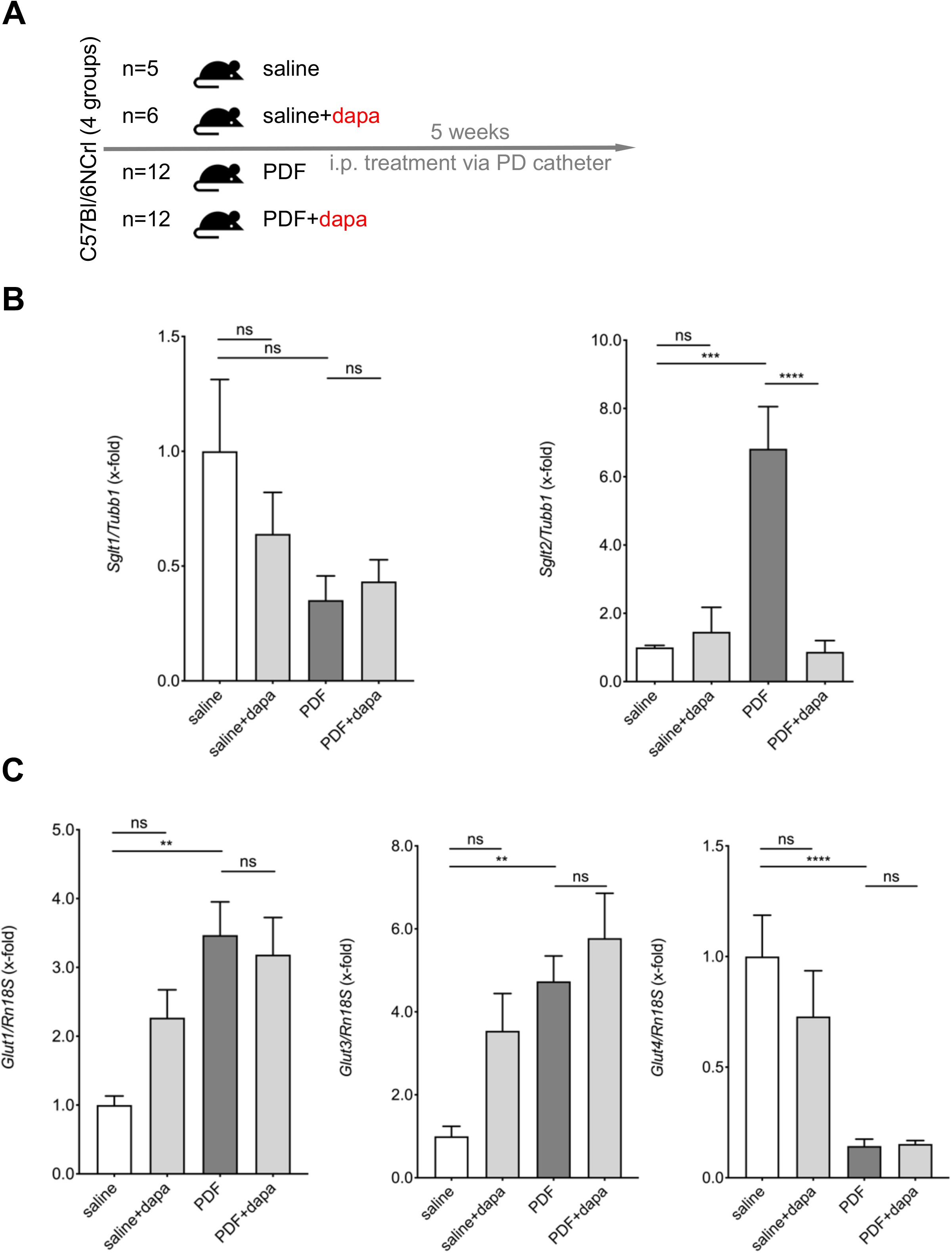
PDF-mediated regulation of peritoneal SGLT and GLUT *in vivo*. (A) Schematic of the study design. C57Bl/6N mice were subjected to 5 weeks of daily treatment with either saline or high glucose (4.25%)-containing PD fluid (PDF) ± dapagliflozin (1 mg/kg body weight). (B) Murine peritoneal membrane mRNA expression of *Sglt1* and *Sglt2*. Expression was normalized to β-tubulin (Tubb1). ns, not significant; *** p<0.001, **** p<0.0001 for Kruskal-Wallis test. (C) Murine peritoneal membrane mRNA expression of *Glut1, Glut3* and *Glut4*. Expression was normalized to *Rn18s*. ns, not significant; ** p<0.01, **** p<0.0001 for Kruskal-Wallis test.

We then evaluated the peritoneal transcriptional expression of SGLT2, SGLT1 and several GLUTs known to be expressed in the peritoneum. We found a strong upregulation of SGLT2 expression in mice receiving high glucose PDF, whereas SGLT1 expression was unaltered (**Figure 2b**). Most notably, pharmacological inhibition of SGLT2 with dapagliflozin completely abrogated PD-induced upregulation of SGLT2. Glucose transporters demonstrated differential regulation, GLUT1 and 3 being upregulated and GLUT4 down-regulated, respectively, in response to chronic exposure to PDF. This regulation was unaffected by dapagliflozin (**Figure 2c**).

In summary, we demonstrated abrogation of PDF-induced SGLT2 transcriptional upregulation by intraperitoneal application of dapagliflozin.

### Peritoneal fibrosis and ultrafiltration failure are ameliorated by dapagliflozin

Having demonstrated a) the existence of SGLT2 at the murine and human peritoneum, b) differential regulatory effects of a high glucose environment on peritoneal glucose transporter expression and c) the effect of pharmacological intervention on expression of SGLT2, we wanted to further evaluate effects of pharmacological SGLT2 inhibition on the development of structural and functional changes in the peritoneal membrane. As expected, pronounced submesothelial thickening and fibrosis developed after a 5 week exposure to PDF (**Figures 3a-b**), accompanied by increased TGF-β levels in effluent (**Figure 3c**). Most importantly, ultrafiltration (UF) decreased, as evaluated after a 120 min intraperitoneal dwell of 4.25% glucose PDF (**Figure 3d**). All aforementioned changes were substantially mitigated by pharmacological SGLT2 inhibition with dapagliflozin. It should be noted, however, that there was a trend towards high glucose-independent structural pro-fibrotic changes in animals receiving saline+dapagliflozin.

**Figure 3:**
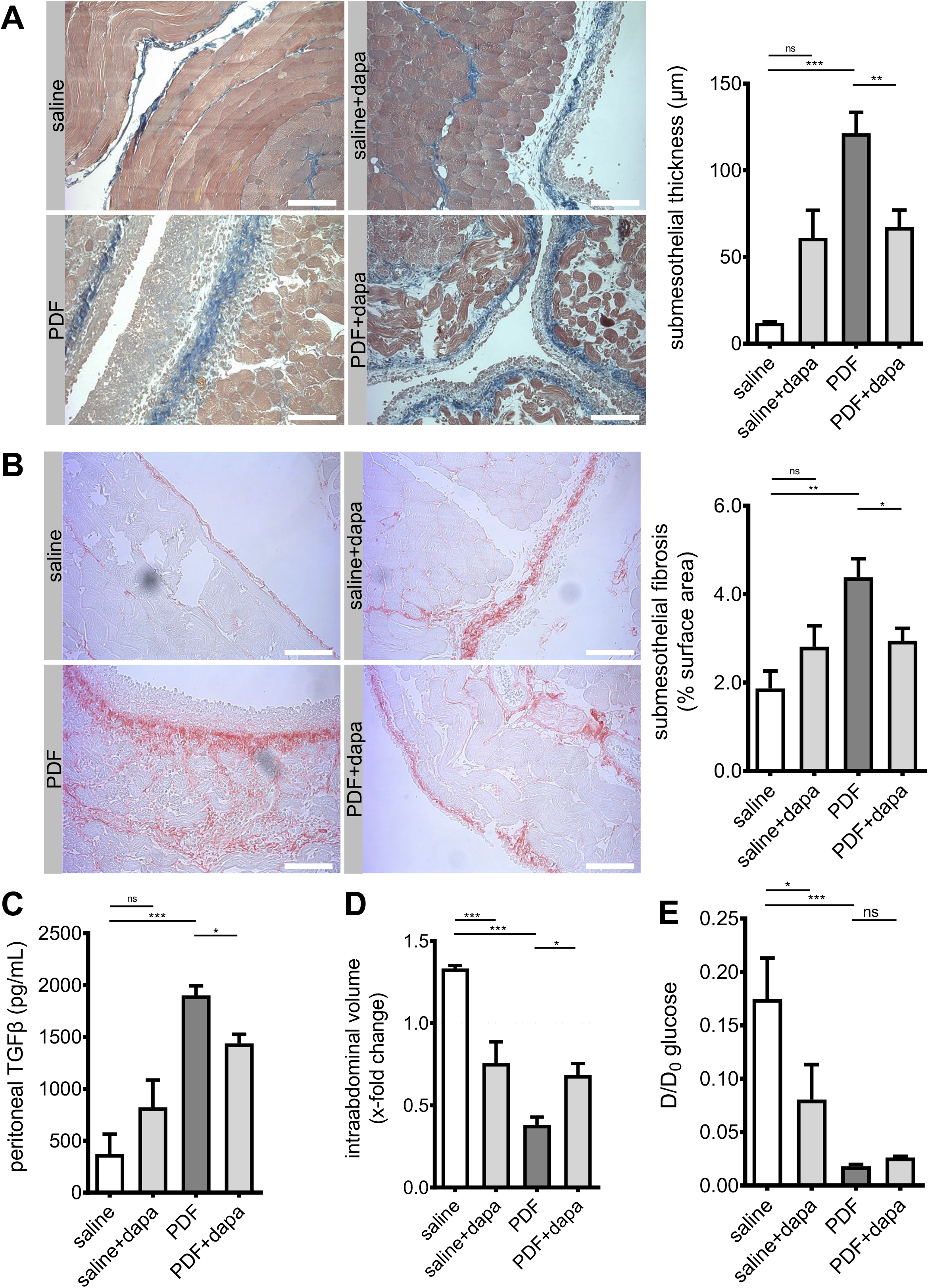
Amelioration of peritoneal fibrosis and ultrafiltration failure by dapagliflozin. (A) Representative images and quantification of Masson’s trichrome staining of murine peritoneum in animals treated with saline, saline+dapagliflozin, peritoneal dialysis fluid (PDF) and PDF+dapagliflozin, respectively. Scale bar=100 μm; ** p<0.01, *** p<0.001 for ANOVA. (B) Visualization of collagen I and III as surrogate for submesothelial fibrosis. Representative images of Picrosirius red staining of murine peritoneum. Scale bar=100 μm. Quantification of percentage of submesothelial fibrosis. * p<0.05, ** p<0.01 for ANOVA. (C) Quantification of peritoneal effluent TGF-β as analyzed by ELISA. * p<0.05, *** p<0.001 for ANOVA. (D) Quantification of ultrafiltration capacity as analyzed by intraabdominal volume after a 120 min challenge with high glucose (4.25%) PDF after 5 weeks of respective treatment conditions. Values >1.0 indicate ultrafiltration, whereas values <1.0 indicate net fluid absorption; * p<0.05, *** p<0.001 for ANOVA. (E) Analysis of glucose transporter status by peritoneal equilibration testing at time points 0 and 120 min, respectively. D and D_0_ denote peritoneal effluent glucose concentrations at time points 120 min and 0, respectively; * p<0.05, *** p<0.001 for Kruskal-Wallis test.

Functionally, dapagliflozin decreased UF capacity in the absence of a high glucose environment (**Figure 3d**), which is consistent with findings from peritoneal equilibration testing (PET), showing that both dapagliflozin and PDF led to a significant decrease of D/D_0_ glucose ratio (**Figure 3e**). The D/D_0_ ratio measures the amount of glucose in dialysate after a 120 min dwell of PDF compared to time 0. The decrease of this ratio indicates a faster reabsorption of glucose, suggesting an acceleration of glucose transport across the peritoneal membrane. This effect of dapagliflozin was specific for glucose, since we noted no changes between PDF-treated animals treated with and without dapagliflozin for other solute transport characteristics: Dialysate-to-plasma ratios (D/P) as well as mass transfer area coefficients (MTAC) for creatinine and urea were similar across all treatment groups (**Supplementary Figure 2**).

In summary, we demonstrate that dapagliflozin reduced peritoneal fibrotic changes, resulting in amelioration of PDF-induced ultrafiltration failure.

### Dapagliflozin reduces submesothelial microvessel density in non-VEGF-dependent manner

As peritoneal transport is influenced by angiogenesis, which is upregulated in response to PDF,[11] we next evaluated microvessel density in CD31-stained sections of murine peritoneum. As expected, PDF-treated animals demonstrated a substantial increase of CD31 positive cells in an area 400 μm below the mesothelial cell layer (**Figure 4a**). Automated counting of microvessels in the submesothelial zone confirmed a significant increase in vessel density (**Figures 4a-b**). Dapagliflozin-treated animals demonstrated reduced microvessel density (p=0.06). Of note, while PDF-treated animals showed a significant increase in vascular endothelial growth factor A (VEGF-A) levels in peritoneal effluents, dapagliflozin-treated animals had similar levels of VEGF-A, suggesting that dapagliflozin-mediated reduction of angiogenesis was independent of VEGF-A (**Figure 4c**).

**Figure 4:**
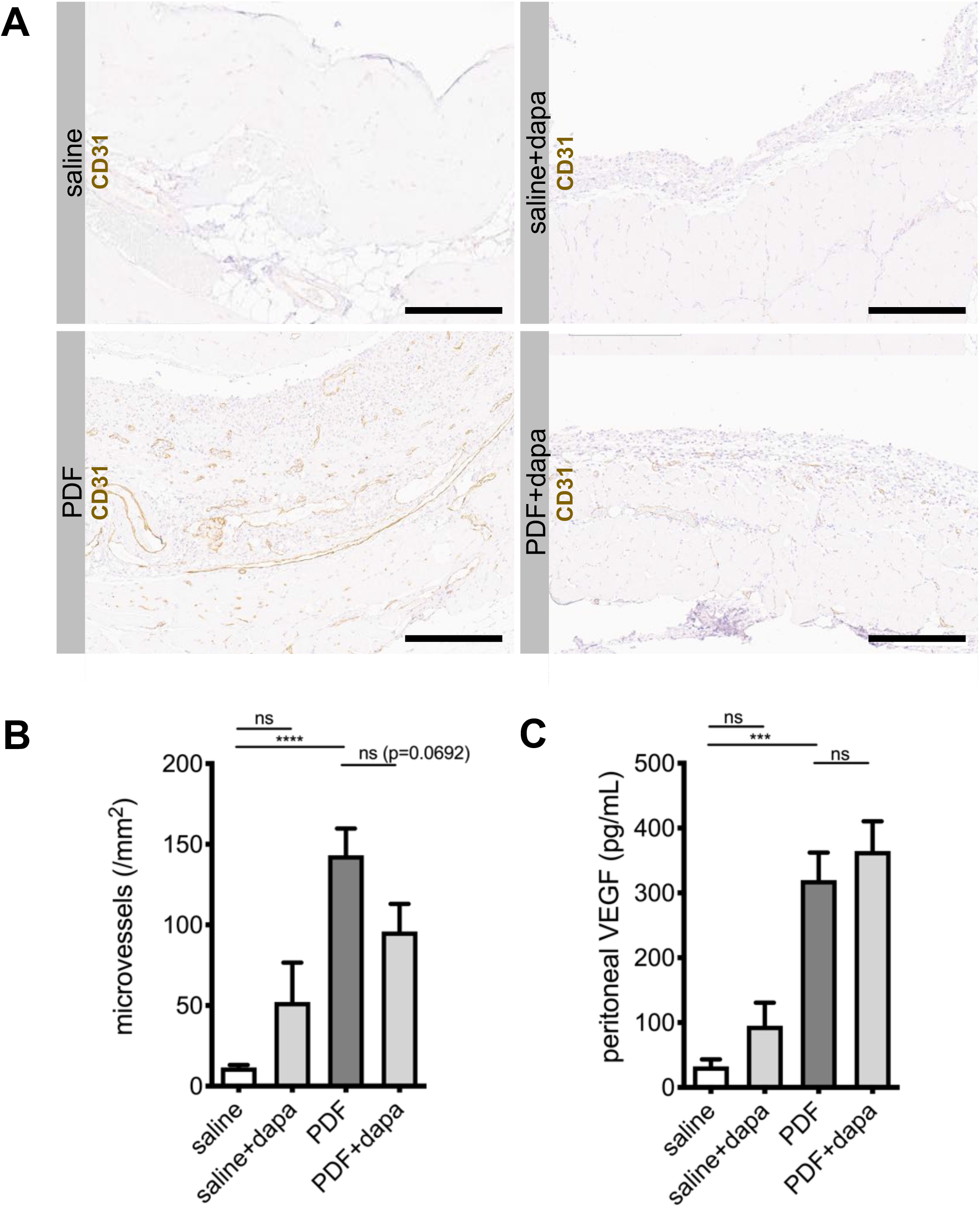
Dapagliflozin reduces submesothelial microvessel density. (A) Representative images of immunohistochemistry staining against CD31 in murine peritoneum in animals treated with saline, saline+dapagliflozin, PDF and PDF+dapagliflozin, respectively. Scale bar=200 μm. (B) Quantification of microvessel density within an area reaching 400 μm below the mesothelial cell layer using Aperio Image Scope microvessel algorithm v1; **** p<0.0001 for ANOVA. (C) Quantification of peritoneal effluent levels for vascular endothelial growth factor A (VEGF-A); *** p<0.001 for ANOVA.

### Dapagliflozin modulates intraperitoneal inflammatory response

After demonstrating ameliorating effects of SGLT2 inhibition on development of peritoneal fibrosis, angiogenesis and UF failure in a high glucose milieu we were interested to evaluate its effects on intraperitoneal inflammation. We therefore analyzed the composition of intraperitoneal cell influx in effluents obtained after a 120 min dwell of PDF across all groups. Consistent with previous findings from our group,[4] chronic PDF exposure led to a significant increase in peritoneal cell count, predominantly leukocytes. Significantly different changes were noted for T cells, B cells, polymorphonuclear neutrophils (PMN) and macrophages (**Figure 5a**). While dapagliflozin had no effect on T and B cell composition, we noted a significantly reduced amount of PMN and an increase in macrophages beyond the PDF-mediated level. Concurrently, intraperitoneal cytokine levels measured in effluents by ELISA demonstrated increases of pro-inflammatory markers IL-6, TNF-α and MCP-1 after PDF exposure (**Figure 5b**). Interferon-□ and anti-inflammatory interleukin-10 also increased in effluents of PDF-exposed mice compared with saline-treated controls, but there were no significant differences in PDF-exposed animals treated with or without dapagliflozin. Again, similarly with pro-fibrotic changes, there was a non-significant trend towards high glucose-independent increase of pro-inflammatory mediators MCP-1 and TNF-α in animals receiving saline and dapagliflozin (**Figure 5b**).

**Figure 5:**
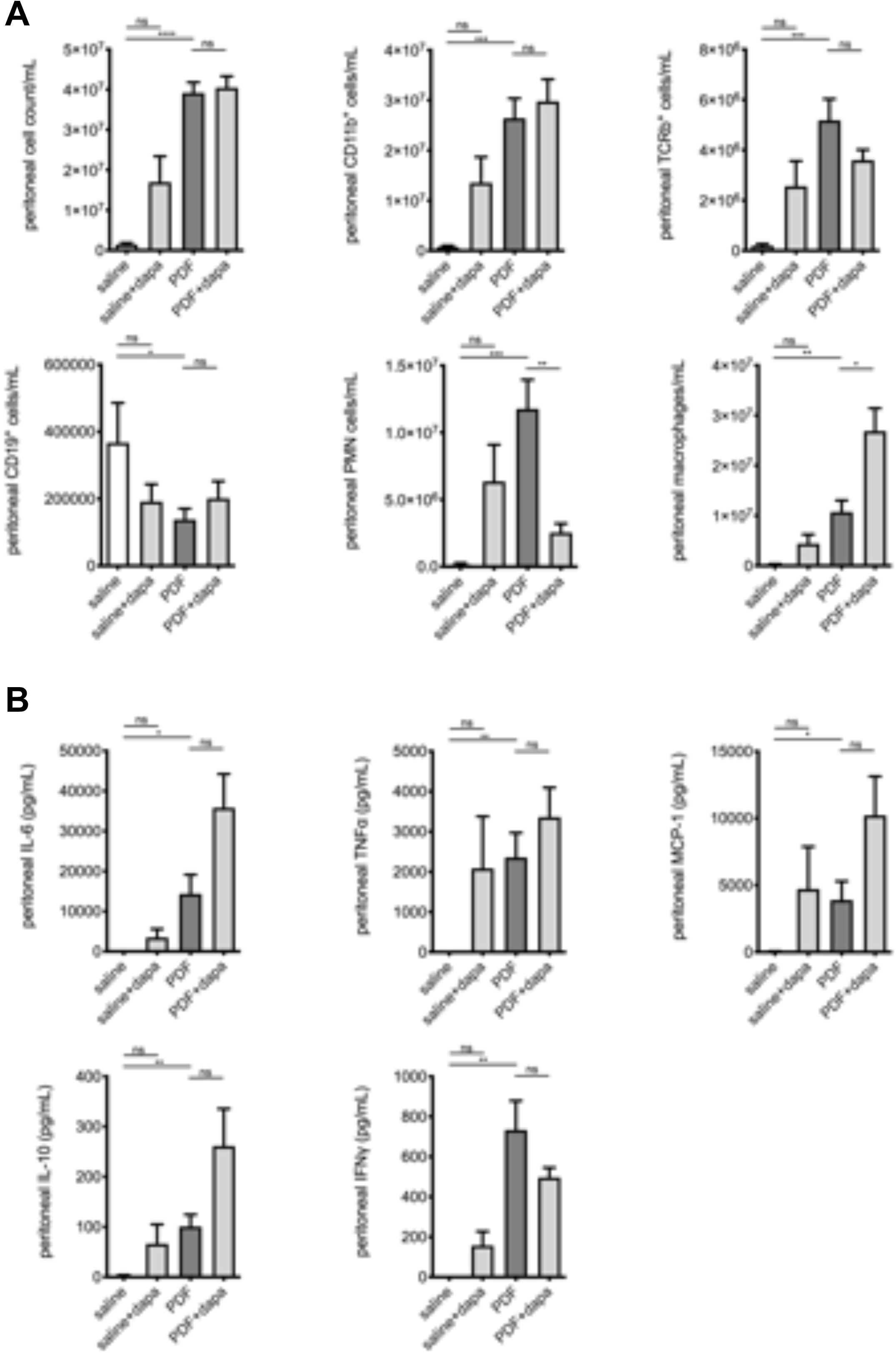
Modulation of intraperitoneal inflammatory response by dapagliflozin. (A) Quantification of inflammatory cell recruitment of total cells, CD11b^+^ cells (leukocytes), αβTCR^+^ cells (T cells), CD19^+^ cells (B cells), polymorphonuclear neutrophils (PMN) and macrophages to the peritoneal cavity, as measured by flow cytometry; * p<0.05, ** p<0.01, *** p<0.001, **** p<0.0001 for Kruskal-Wallis test or ANOVA, as applicable. (B) Quantification of peritoneal effluent levels of pro-inflammatory cytokines interleukin-6 (IL-6), tumor necrosis factor-α (TNF-α) and monocyte chemoattractant protein-1 (MCP-1), anti-inflammatory cytokines IL-10 and interferon-□ (IFN-□); * p<0.05, ** p<0.01, *** p<0.001 for Kruskal-Wallis test or ANOVA, as applicable.

### Dapagliflozin abrogates pro-inflammatory signaling in murine and human peritoneal mesothelial cells and exerts glucose-independent anti-inflammatory effects on murine peritoneal macrophages

As SGLT2 inhibition significantly ameliorated *in vivo* fibrotic and functional changes and had equivocal effects on inflammatory response, we wanted to further analyze the effects of dapagliflozin on mesothelial cells and macrophages *in vitro*. In murine omentum-derived mesothelial cells, only SGLT2, but not SGLT1 transcription was upregulated in response to dapagliflozin in a high glucose environment (**Figure 5a**).

Pharmacological inhibition of SGLT2 decreased both glucose consumption and uptake in HPMC. [15] We therefore asked whether dapagliflozin decreases intracellular glucose content in murine mesothelial cells and macrophages cultured under high glucose conditions. Expectedly, glucose concentration in lysates of both MPMC and macrophages significantly increased in high glucose conditions (**Figure 6b)**. Dapagliflozin reduced glucose uptake in a dose-dependent manner both under normal and high glucose conditions in MPMC. In contrast, in murine macrophages, dapagliflozin affected glucose uptake only in a high glucose milieu. It should be noted that high glucose-induced increase of intracellular glucose concentration was only partially reduced and not completely normalized by dapagliflozin in either cell type.

**Figure 6:**
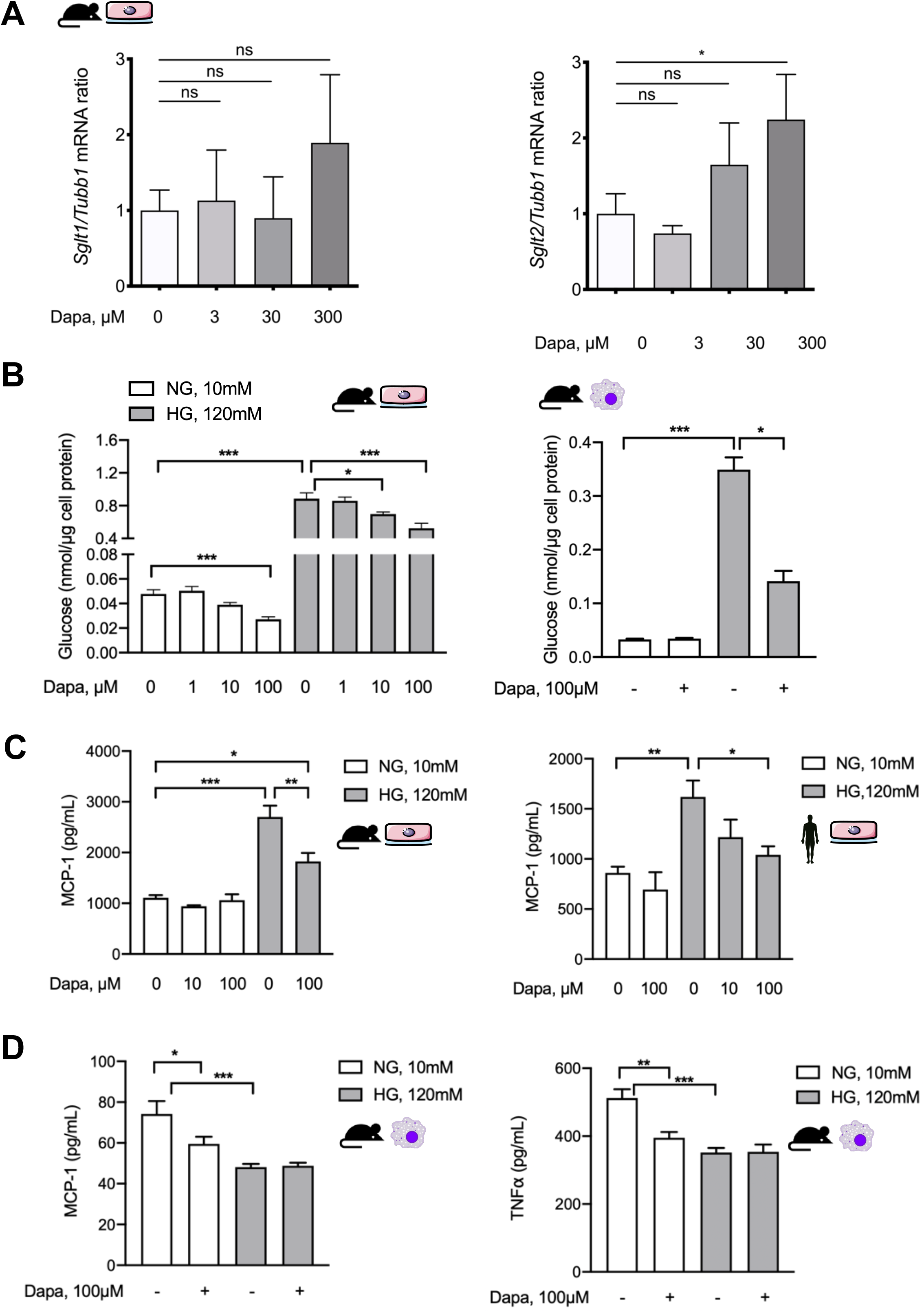
Dapagliflozin abrogates pro-inflammatory signaling *in vitro* in mesothelial cells and exerts anti-inflammatory effects in macrophages. (A) Quantification of mRNA expression of *Sglt1* and *Sglt2* in MPMC in response to high glucose conditions with or without additional dapagliflozin treatment in ascending concentrations. * p<0.05 for Kruskal-Wallis test. (B) Quantification of glucose concentration in lysates of MPMC (left) and murine macrophages (right) as a surrogate of cellular glucose uptake under either normal or high glucose conditions with or without additional dapagliflozin treatment for 48 h in different concentrations. NG, normal glucose (10 mM); HG, high glucose (120 mM); * p<0.05, *** p<0.001 for Kruskal-Wallis test. (C) Quantification of MCP-1 in conditioned medium from MPMC and HPMC culture under either normal or high glucose conditions for 24 h with or without additional dapagliflozin treatment in different concentrations; * p<0.05, ** p<0.01, *** p<0.001 for Kruskal-Wallis test. (D) Quantification of MCP-1 and TNF-α in conditioned medium from murine macrophages cultured for 24 h under either normal or high glucose conditions with or without additional dapagliflozin treatment; * p<0.05, ** p<0.01, *** p<0.001 for Kruskal-Wallis test.

The effect of dapagliflozin on high glucose-induced MCP-1 production in mesothelial cells and macrophages was analyzed next. Both in murine (MPMC) and human peritoneal mesothelial cells (HPMC) dapagliflozin reduced MCP-1 release in a high glucose milieu, while it had no effect in normal glucose conditions (**Figure 6c**). Consistent with our previous findings, mesothelial cells increased MCP-1 production in response to high glucose, while murine macrophages produced less, possibly reflecting a shift to M2 polarization under high glucose conditions.[13] Similarly, dapagliflozin administered in normal glucose conditions reduced MCP-1 release but had no further effect in high glucose conditions. These effects were also observed for TNF-α production (**Figure 6d**).

Given our observation of increased peritoneal macrophages in peritoneal effluents in PDF+dapagliflozin-treated animals, we evaluated dapagliflozin action in murine macrophages under similar normal vs. high glucose conditions with an additional pro-inflammatory stimulus. To this end, we used an experimental setup where the cells were first cultured for 48 h under normal or high glucose conditions with or without dapagliflozin and thereafter additionally stimulated with 100 ng/mL LPS for 8h (**Figure 7**). In line with previous observations, increased macrophage production of pro-inflammatory mediators MCP-1 (**Figure 7a**), TNF-α (**Figure 7b**) and IL-6 (**Figure 7c**) upon LPS stimulation was toned down in a high glucose compared to a normal glucose environment. This signature is well-known for M2 macrophages, which make up a considerably larger fraction of macrophages in glucose-mediated PM damage and for which we have previously demonstrated glucose to be the decisive driver of this M1-to-M2 switch.[13] Dapagliflozin decreased pro-inflammatory response only under normal glucose conditions, but not in high glucose conditions (**Figure 7c**), thereby tying in with our *in vivo* observations in PDF-treated mice.

**Figure 7:**
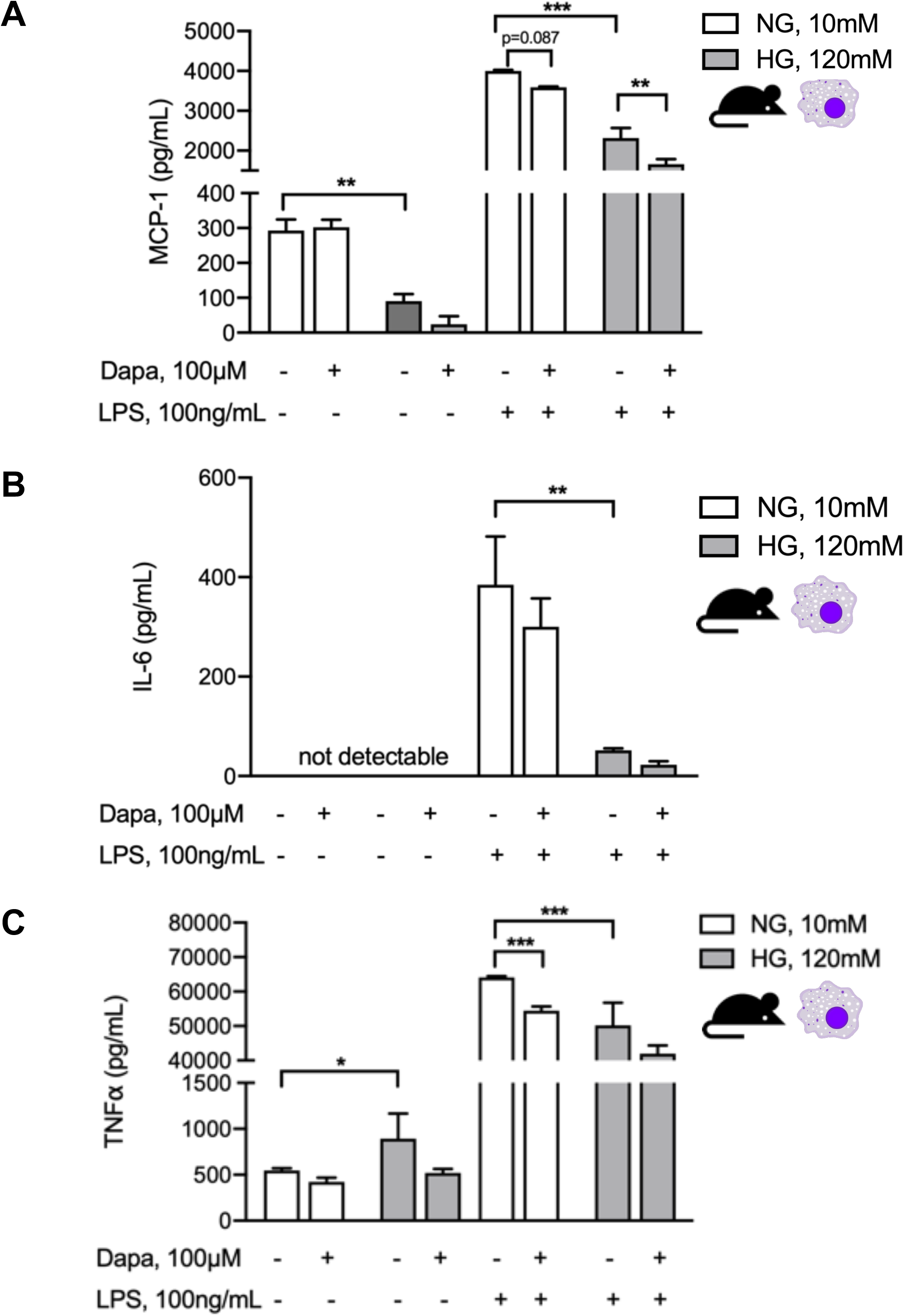
Dapagliflozin abrogates pro-inflammatory stimuli in murine macrophage culture. Quantification of MCP-1 (A), IL-6 (B) and TNF-α (C) in supernatants of murine macrophages under either normal or high glucose conditions with or without additional dapagliflozin treatment for 48 h and with or without subsequent stimulation with LPS for 8 h. ** p<0.01, *** p<0.001 for Kruskal-Wallis test.

## DISCUSSION

Glucose has been implicated as a major mechanism of peritoneal membrane pathophysiology in PD.[16,17] The chronic peritoneal glucose exposure induces significant systemic sequelae. We have previously shown that daily dialytic glucose exposure is associated with vascular complement and TGF-β activation and closely correlated with the degree of vasculopathy.[18] The major route of glucose uptake into mammalian cells is through either facilitative glucose transporters (GLUT)[19] or sodium-driven glucose symporters (SGLT).[20] These glucose transporters are cell-specifically expressed and have specialized glucose-sensing properties, which contribute to glycolysis and related cellular functions.[21] However, information about glucose uptake into mesothelial cells and its regulation is scarce. From cell culture studies in HPMC we know that GLUT mRNA expression and glucose uptake are induced by high ambient glucose concentration as well as by proinflammatory cytokines.[22] In addition, it has been known for over three decades that protein kinase C (PKC) activation rapidly initiates glucose uptake into cells and may phosphorylate GLUT1.[23] Given that we have previously shown that classical PKC isoform α in mesothelial cells is responsible for glucose-mediated peritoneal membrane damage,[4] it is interesting that the phosphorylation site for conventional and novel PKCs was only recently identified in GLUT1,[24] highlighting the importance of cellular glucose transport. In addition to GLUT, the expression of SGLT1 and SGLT2 in mesothelial cells cultured *in vitro* has already been reported.[6,15] Despite the recent surge of information on SGLT2, however, it is less clear whether this protein might represent a viable target for influencing peritoneal health.

In the present study, we demonstrate that both SGLT1 and SGLT2 are expressed in the peritoneal membrane in mice and humans. In mice, chronic exposure of the peritoneal membrane to a high glucose milieu in PDF regulated the expression of glucose transporters such as GLUT1, GLUT3, GLUT4 and SGLT2. We show for the first time that SGLT2 inhibition via intraperitoneal application of dapagliflozin ameliorates structural and functional changes in PDF-induced peritoneal fibrosis. Dapagliflozin/PDF treatment reduced peritoneal thickening and fibrosis and improved ultrafiltration compared to animals treated with PDF alone. These changes are in keeping with evidence from other organs where SGLT2 inhibition has been associated with antifibrotic effects, most prominently in the kidney but also in the heart and the liver. For example, dapagliflozin promoted antifibrotic effects in a type 1 diabetic kidney disease model by ameliorating O-GlcNAcylation and reducing tubular hypoxia,[25] while others have found a downregulation in the Stat1/TGF-β pathway as well as decreased epithelial-to-mesenchymal transition.[26] Importantly, beneficial effects of SGLT2 inhibition were also demonstrated in non-diabetic kidney disease such as hypertensive nephropathy and were attributed to anti-inflammatory effects. [27] Similarly, anti-fibrotic effects of SGLT2 inhibition were found in liver[9] and heart,[10] where administration of dapagliflozin reduced cardiac fibrosis by stimulating M2 macrophages and inhibiting myofibroblast differentiation. [28] Microvessel density in the first 400 μm below mesothelial cell layers, representing the penetration level of PD fluids,[29] was increased in PDF-treated animals and additional dapagliflozin treatment mitigated this PDF-induced increase. In PD patients, within a few months after PD start, glucose-containing PDF induces an increase of peritoneal microvessel density, which is associated with peritoneal membrane transport function at baseline and during PD.[11] Thus, reduced microvessel density may have contributed to mitigation of PD-induced UF loss. However, since the D/D_0_ glucose ratio was not improved, the substantially reduced fibrosis may have contributed to the mitigated UF loss by improving osmotic conductance to glucose, a significant determinant of UF in PD. [30] The effluent cytokine analyses suggest a VEGF-independent mechanism of reduced peritoneal vascularization by SGLT2 inhibition and argue in favor of pathways such as modulation of angiopoietin 1/2,[31] but determination of tissue cytokine abundance may be more sensitive and valid.

As mentioned above, we observed some glucose-independent detrimental side effects of dapagliflozin. The dose of 1 mg/kg of dapagliflozin used in our experiments has been shown as safe and well tolerated in mice if given systemically up to 12 weeks.[32,33] In order to achieve effective concentrations of dapagliflozin in the peritoneal cavity, we used an intraperitoneal way of administration. It should be noted that despite local application of dapagliflozin, systemic action was observed, as reflected by presence of glucosuria in 24 h urine collections of mice treated with the SGLT2 inhibitor. This suggests uptake by peritoneal blood capillaries or by lymphatics, which is not surprising given the low molecular weight of dapagliflozin. However, we cannot fully exclude intraperitoneal accumulation of dapagliflozin leading to increased local concentration, which might possibly result in local toxic effects. It would be interesting to test whether systemic application of dapagliflozin will still have a protective effect at the PM without possible detrimental glucose-independent side effects observed by local application. In the setting of a high glucose environment, however, we saw significant benefits of additional dapagliflozin application.

Despite the marked reduction of peritoneal fibrotic changes and microvessel density with SGLT2 inhibition in our PDF exposure model, dapagliflozin action on glucose-mediated peritoneal inflammatory response was more complex. While cell influx to the peritoneal cavity was unchanged with regard to T and B cells, we noted a significant reduction of PMN and increase in peritoneal macrophages. At the same time, dapagliflozin administered in the absence of a high glucose milieu showed a trend towards PM thickening (p=0.10), whereas increases in proinflammatory cytokines such as IL-6, TNF-α and MCP-1, and anti-inflammatory cytokines such as IL-10 and IFN-□ were not statistically significant. Given these equivocal *in vivo* results, we tried to pinpoint the influence of a high glucose milieu and SGLT2 inhibition on inflammatory responses of mesothelial cells and macrophages *in vitro*. Interestingly, mesothelial cells demonstrated dapagliflozin-mediated reduction of MCP-1 only in the presence of a high glucose milieu, whereas in macrophages this reductive effect was only noticeable in normal glucose conditions. There is an accumulating body of evidence that macrophages are important targets for anti-inflammatory effects mediated by SGLT2 inhibition. For example, SGLT2 inhibition prevented inflammation via inhibition of macrophage accumulation and MCP-1 expression. [34] In another study, empagliflozin inhibited MCP1 and TGF-β gene expression in an experimental model of diabetic nephropathy,[35] while others have found that SGLT2 inhibition reduced levels of MCP-1, IL-6 and TNF-α in aortic plaques and adipose tissue,[36] as well as nuclear factor κB and IL6 levels in renal tissues.[37] Also, respective effects have been attributed to polarization of macrophages towards an M2 phenotype.[38] Here, we studied the effects of SGLT2 inhibition on macrophages challenged with an inflammatory stimulus in both normal and high glucose milieus. To this end, we stimulated macrophages with LPS after treating them with either normal or high glucose conditions in presence or absence of dapagliflozin. We observed that when macrophages were not challenged by LPS, dapagliflozin had no effect on proinflammatory marker release, whereas MCP-1 and TNF-α were significantly reduced by SGLT2 inhibition in the presence of an inflammatory stimulus such as LPS. This effect was seen in cells cultured under normal glucose conditions (**Figure 6d**), suggesting that SGLT2 inhibition can shift macrophages to M2 polarization independently from glucose. This observation is in line with a recently published novel mechanism of SGLT2 inhibitors-mediated M2 polarization through a glucose-independent reactive oxygen and nitrogen species-dependent STAT3-mediated pathway.[28] In line with this, when macrophages were cultured under high glucose condition and therefore already shifted to M2 prior to LPS stimulation,[13] anti-inflammatory effects of dapagliflozin were still observed albeit less pronounced and reached statistical significance for MCP-1 only (**Figure 7**).

In summary, our data demonstrate the presence of SGLT2 in the murine and human peritoneum, its regulation by glucose in mice and beneficial effects of its inhibition by dapagliflozin on peritoneal and mesothelial cell health *in vivo* and *in vitro*. Further studies defining the cellular pathways of SGLT2 inhibition influencing peritoneal membrane pathophysiology are warranted in order to understand whether its use in PD patients is a viable treatment option.

## AUTHORSHIP STATEMENT

MSB and NS conceived the research design and had full access to the data; MSB, SR, JN, JDZ, BS, MB, SvV, SS, HH, CPS and NS performed the experiments or were involved in data acquisition; MSB, MB, SvV, CPS and NS analyzed the data; MSB wrote the original version of the manuscript, all authors participated in the review and editing of the manuscript.

## ACKNOWLEDGEMENT

We are grateful to Professor R. Lichtinghagen (Institute of Clinical Chemistry, Hannover Medical School) and Mrs. H. Chlebusch (Department of Nephrology and Hypertension, Hannover Medical School) for technical assistance with serum and urine measurements, to Professor E. Herpel and Mrs. B. Walter (Tissue Bank of the National Center for Tumor Diseases, NCT, Heidelberg and Institute of Pathology, Heidelberg University Hospital) for technical assistance with microvessel density measurements.

---

**Supplementary Figure 1:**
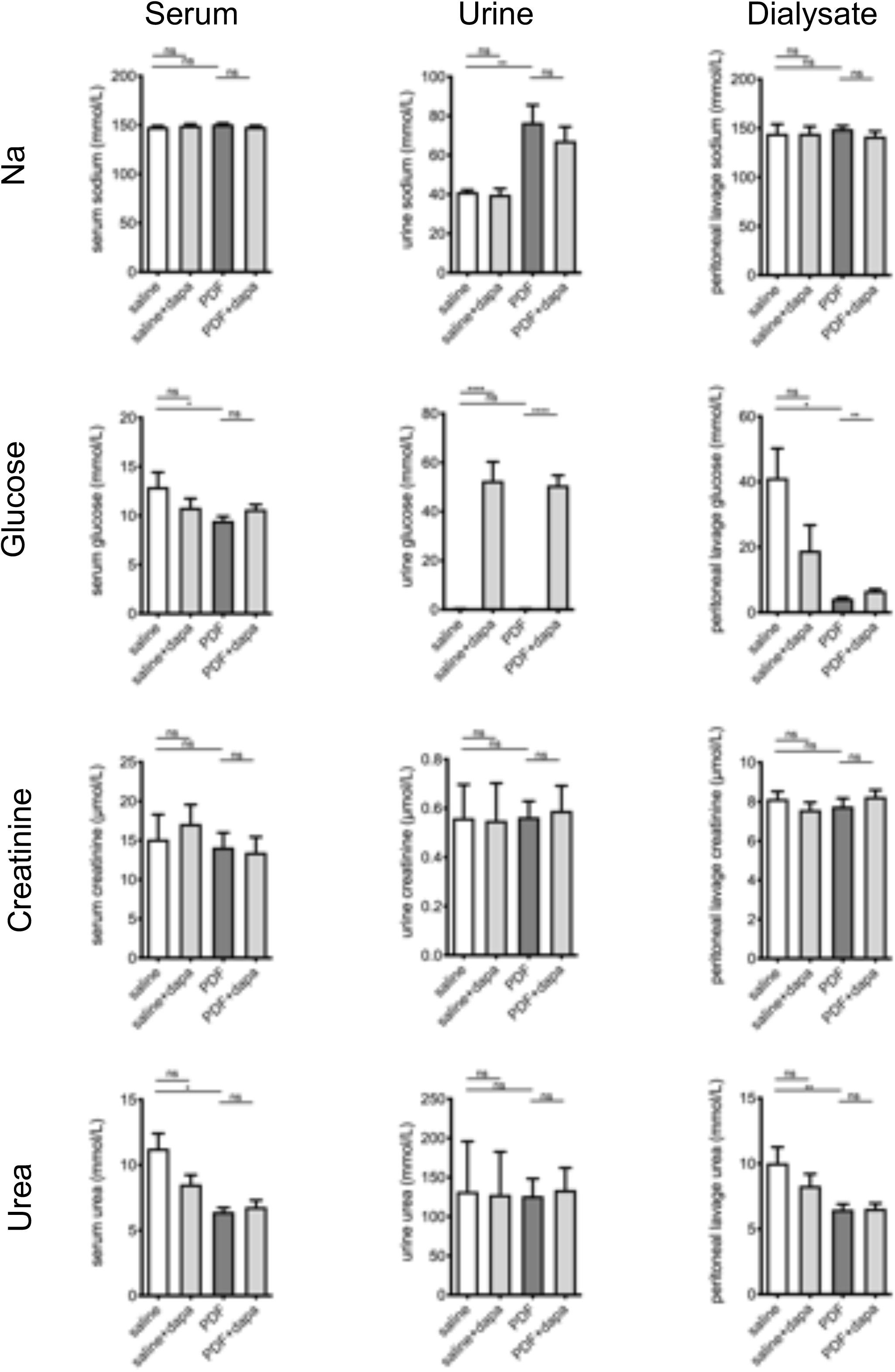
Systemic action of i.p. dapagliflozin. Chemical analysis of serum, urine and peritoneal lavage (dialysate) for sodium (Na), glucose, creatinine and urea, respectively. Presence of glucosuria as measured in 24 h urine collections demonstrates systemic dapagliflozin absorption and action in mice treated with dapagliflozin. ns, not significant; * p<0.05, ** p<0.01, **** p<0.0001 for ANOVA.

**Supplementary Figure 2:**
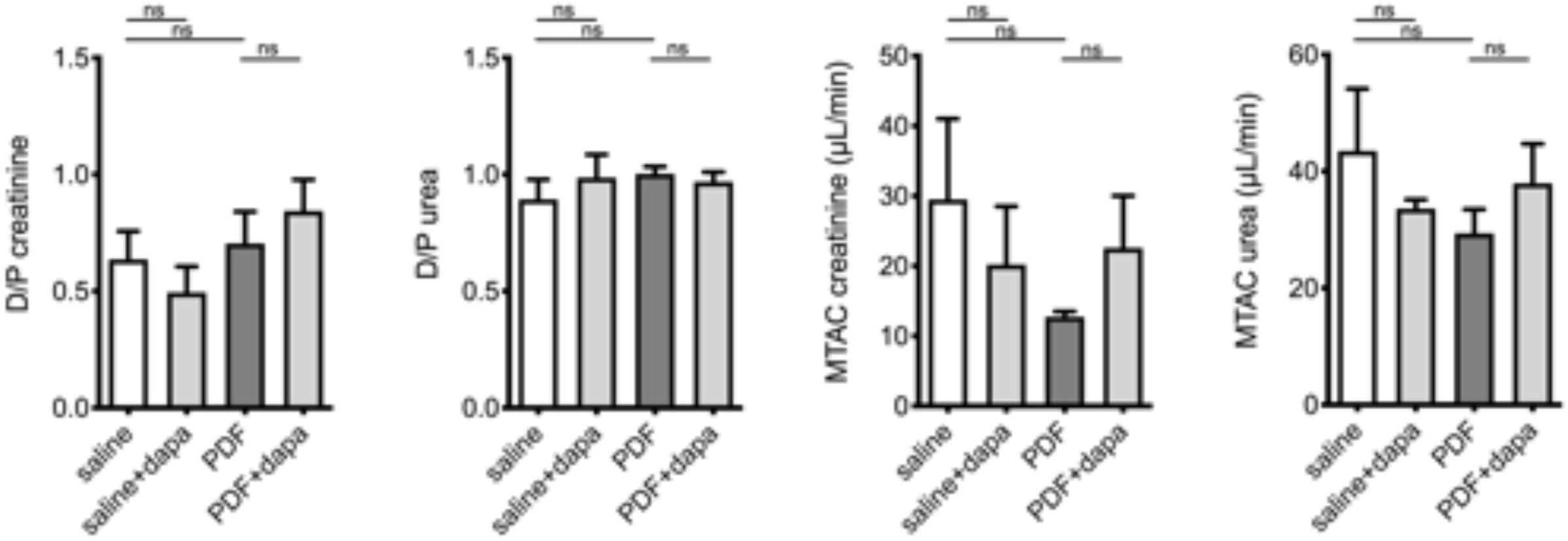
Dapagliflozin does not change peritoneal transport characteristics for uremic solutes. Analysis of dialysate (D)-to-plasma (P) ratios as well as mass transfer area coefficients (MTAC) for creatinine and urea as surrogates of solute transport across the peritoneum. ns, not significant.

